## Supplementary Information for "Smart hybrid microscopy for cell-friendly detection of rare events"

### Supplementary Material

#### 1- Phototoxicity of imaging modalities

To quantify the potential improvement achievable by our smart acquisitions, we performed phototoxicity experiments comparing different imaging modalities. We interleaved the measurements for all modalities on the same sample to reduce bias in the comparisons (Fig S1a), and stained the sample with a Hoechst dye and SYTOX orange to quantify cell death. The Hoechst channel allowed for segmentation of the nuclei, while SYTOX orange overlap with the nuclei regions was used to signal the early stages of cell death. For fluorescence and phase contrast, we took 249 frames with 100 ms exposure at 1 fps for two different fluorescence illumination intensities, and imaged the sample in all channels every 250 seconds for all conditions (Fig. S1b).

To assess the image quality corresponding to the phototoxicity of a certain condition, we quantified signal-to-noise ratio (SNR) and contrast (Fig. S1b). Due to the different characteristics of the images in phase contrast and fluorescence, we opted for a manual labeling approach for mitochondria as foreground and adjacent regions as background that did not show distinctive features in phase contrast. Metrics were calculated by comparing the data in these regions for the first 1000 frames and absolute values are reported in the corresponding panels. SNR was calculated as mean foreground intensity over standard deviation of the background and contrast as mean foreground intensity minus mean background intensity.

In phase contrast and dark conditions, we detected less than 10% cell death across experiments.The results were indistinguishable from one another and similar in both SNR and contrast. On the other hand, using typical fluorescent imaging conditions for MitoTracker (10% laser power), we observed noticeable cell death already at the first measurement timepoint, reaching a half-life time of 8.5 minutes when fitting an exponential decay curve (Table S1). This results in a decay rate ratio of 53 and an excess mortality in fluorescence imaging of 4800 (Eq 1 and 2). Lowering the laser power to 2% reduced the cell death phenotype, still reaching 40% after an hour, with a marked loss in image quality (SNR and contrast).

Taken together, our data suggests that while fluorescence presents a trade-off between image quality and cell death even at low-dose widefield imaging conditions, phase contrast imaging can be performed at high framerates during long acquisitions with minimal cell toxicity and image quality loss.

| Method | Decay Rate 1/h | Half-Life Time |
|---|---|---|
| Fluorescence (10 %) | 4.9 | 8.5 mins |
| Phase Contrast | 0.091 | 7.6 h |
| Dark | 0.090 | 7.7 h |

**Table S1:** Decay rates and half-life times for each condition.

$$\text{Relative decay rate: } \lambda_{fluo}/\lambda_{phase} = 53 \tag{1}$$

$$\text{Relative excess mortality: } \frac{\lambda_{fluo} - \lambda_{dark}}{\lambda_{phase} - \lambda_{dark}} = 4800 \tag{2}$$

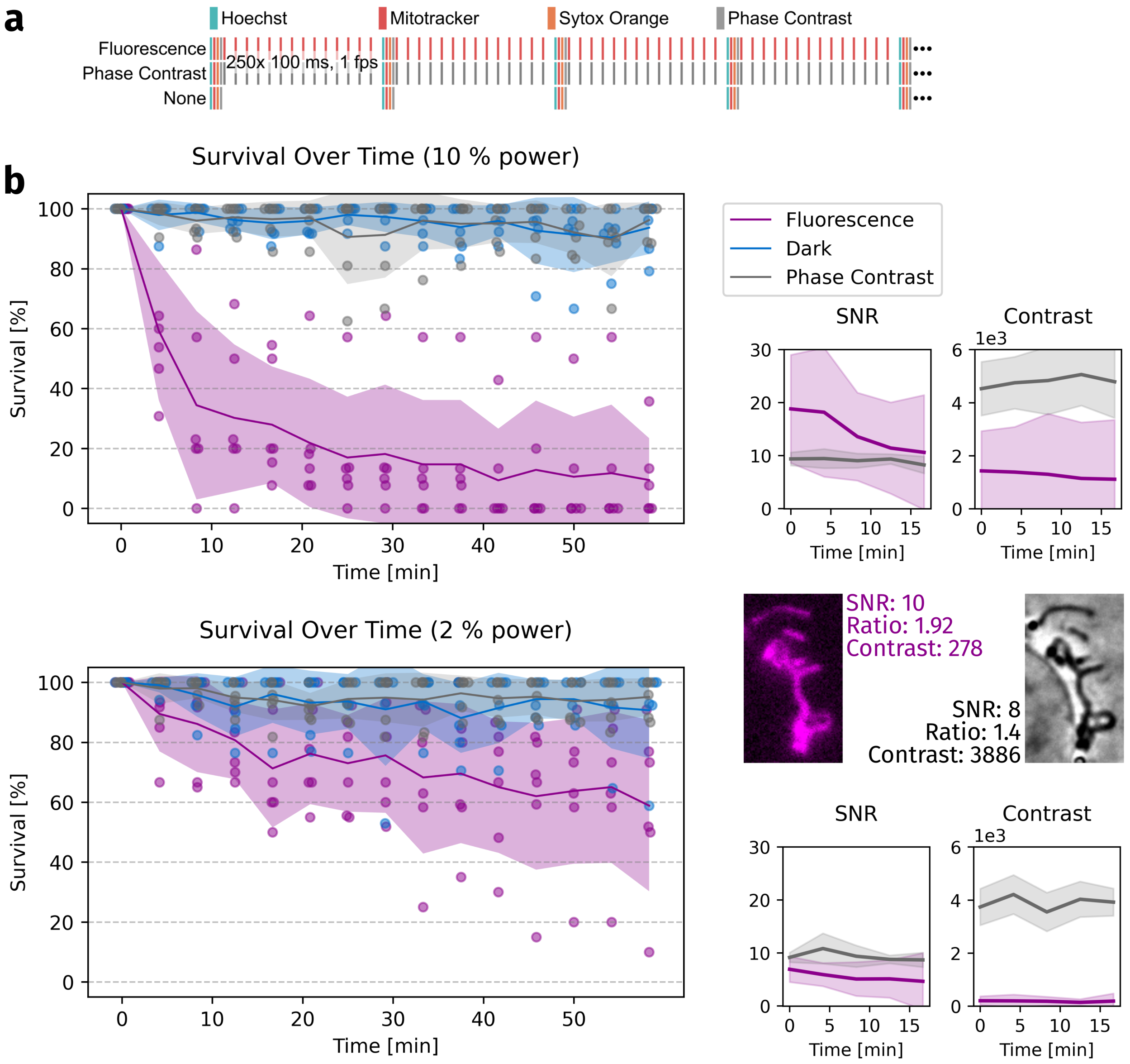


**Fig. S1 | Label-free acquisition cause greatly reduced levels of phototoxicity compared to fluorescence imaging.** **a,** Graphic depiction of imaging setup, vertical lines represent acquired images. **b,** Survival plots for SYTOX toxicity experiment, each point represents an independent FOV (left) and image quality measures in each experiment, including inset examples with measured values (right). Shaded lines represents mean ± sd.

#### 2 - Deep Events data handling

In order to handle the data for this project we have established a database backed data ingestion pipeline. It handles data from different sources (microscopes/softwares) and samples and organizes both image and metadata. Specific to our application is the integration of time lapses and sparse annotations. The balance between negative and positive instances in the training data is important for the training process. As events are randomly distributed, we crop events from the full frame time lapses in space and time. Additionally we add sequences that do not contain events as negatives. Extracted sequences and the corresponding metadata are added to a central database, that can be queried for combinations of data (frame rates, illumination settings etc.). For training, a prompt into the database produces corresponding training and evaluation datasets. Together with the model, we save the metadata for the training (prompt, model parameters etc) and the performance data (scores, prediction histograms etc). To optimize subsequent trainings, we have implemented a manual curation workflow that enables addition of training data to the database when one of the models is used. False positives indicate valuable training data for the model to become more precise for example. As data is added to the database continuously, we have implemented an automatic re-training procedure that trains models on a weekly basis.

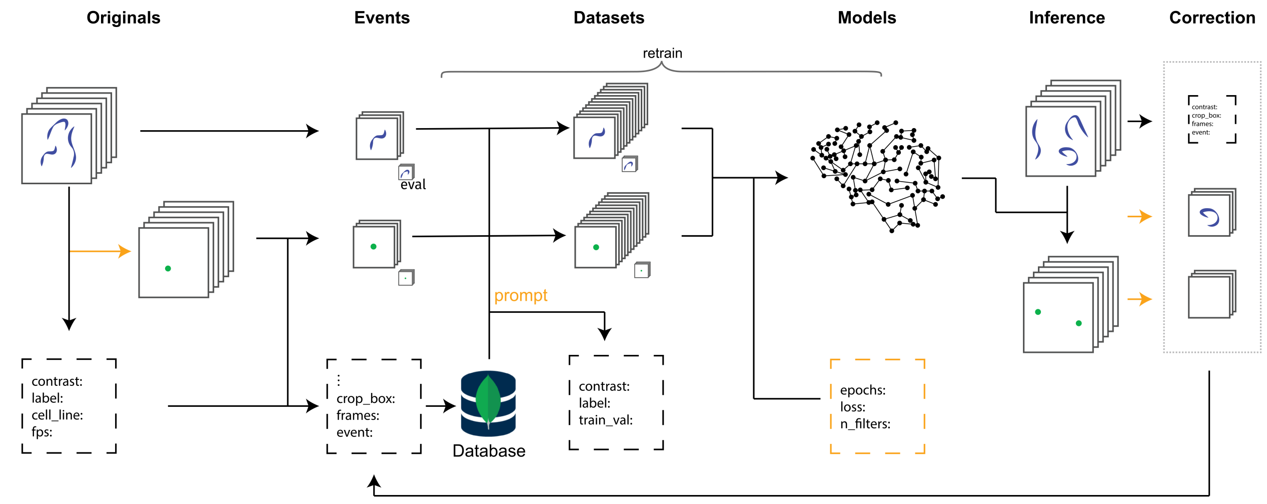


**Fig. S2 | Overview of the data flow for the deep-events package.** Raw microscopy data flows through event extraction, database storage with metadata integration, and dataset generation for model training. A central database enables balanced sampling of positive/negative instances, with automatic retraining procedures ensuring model improvement over time.

#### 3 - Software structure

Commercial microscope software is implemented with the traditional linear approach to multi-dimensional acquisitions (MDA), where a predefined time series of instructions is sent to the microscope for acquisition. Adaptive acquisitions, in the way we have implemented them here, are not supported by this approach. We have therefore worked together with the pymmcore-plus team to implement adaptive acquisitions in the pymmcore-plus framework[1,2]. The surveillance acquisition is sequenced by a slightly adapted MDARunner that uses the MDAEngine to relay MDAEvents to the microscope via the pymmcore (python) bridge to the Micro-Manager core (C++). The microscope (camera) returns the images and an event is published in the events backend in pymmcore-plus. Our on-the-fly analysis pipeline subscribes to this event to be notified for the arrival of new data (Fig S2).

A component called analyser handles the data ingestion, stacking time frames for the model and filtering data from other channels. It implements preprocessing of the frames and runs inference of the model once enough frames have arrived. The resulting event score map is processed and a final event score for the current time point is sent to an interpreter. The interpreter uses this data to send the decision to activate or deactivate the additional channel(s) to the actuator. It implements memory, allowing advanced settings like a minimal number of frames to be taken in the smart acquisition settings. The actuator stores the additional acquisition settings and changes the active acquisition sequences in the smartRunner. With this the loop is closed and the runner is informed which events to send to the MDAEngine at any given time point.

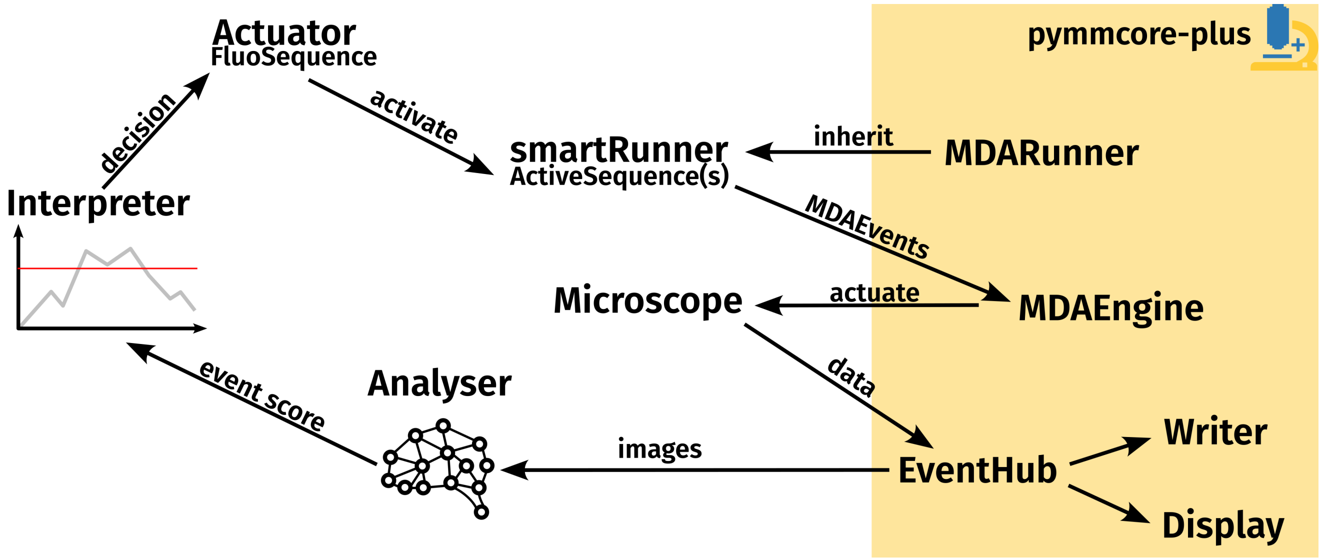


**Fig. S3 | Graphical representation of the software structure.** The software components interact with pynmmcore-plus via efficient event-based communication. Event-scores are calculated from images by the analyser. The interpreter contains long-term temporal context and uses this information to decide on hybrid-EDA activation. Switching betweenmodalities is implemented by the actuator and smartRunner. Pymmcore-plus is used for the basic functionality of microscope control, visualization and data output.

#### 4 - Additional organelle contact events

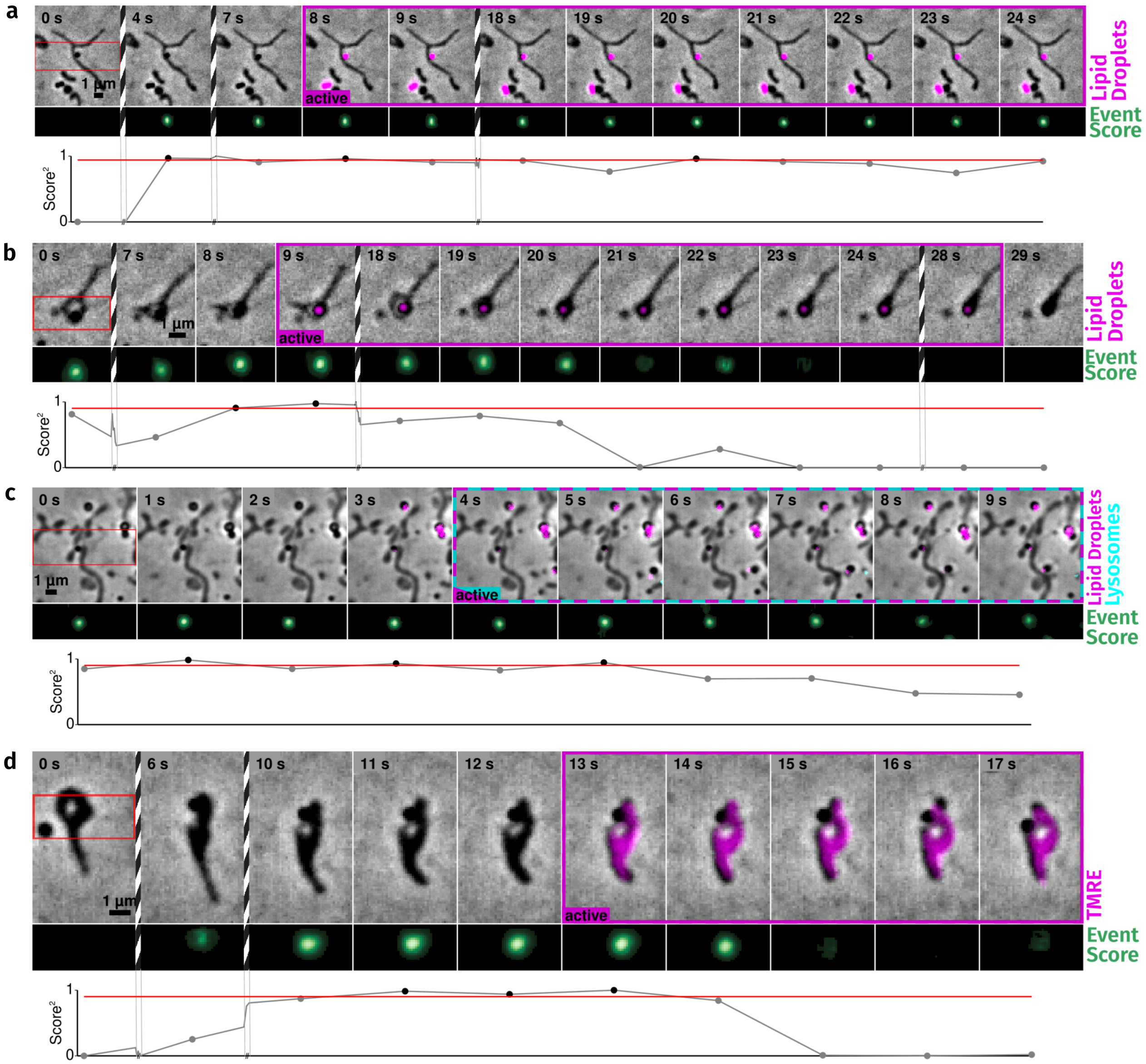


**Fig. S4 | Additional examples of organelle contact events acquired by hybrid-EDA microscopy.** Phase contrast imaging overlaid with fluorescence, as labeled on the right. Below each image time series is the score obtained from the event detection model as an image (crop marked as a red square at time 0) and an integrated plot. The red line indicates the trigger threshold, vertical striped lines indicate cropped timepoints.

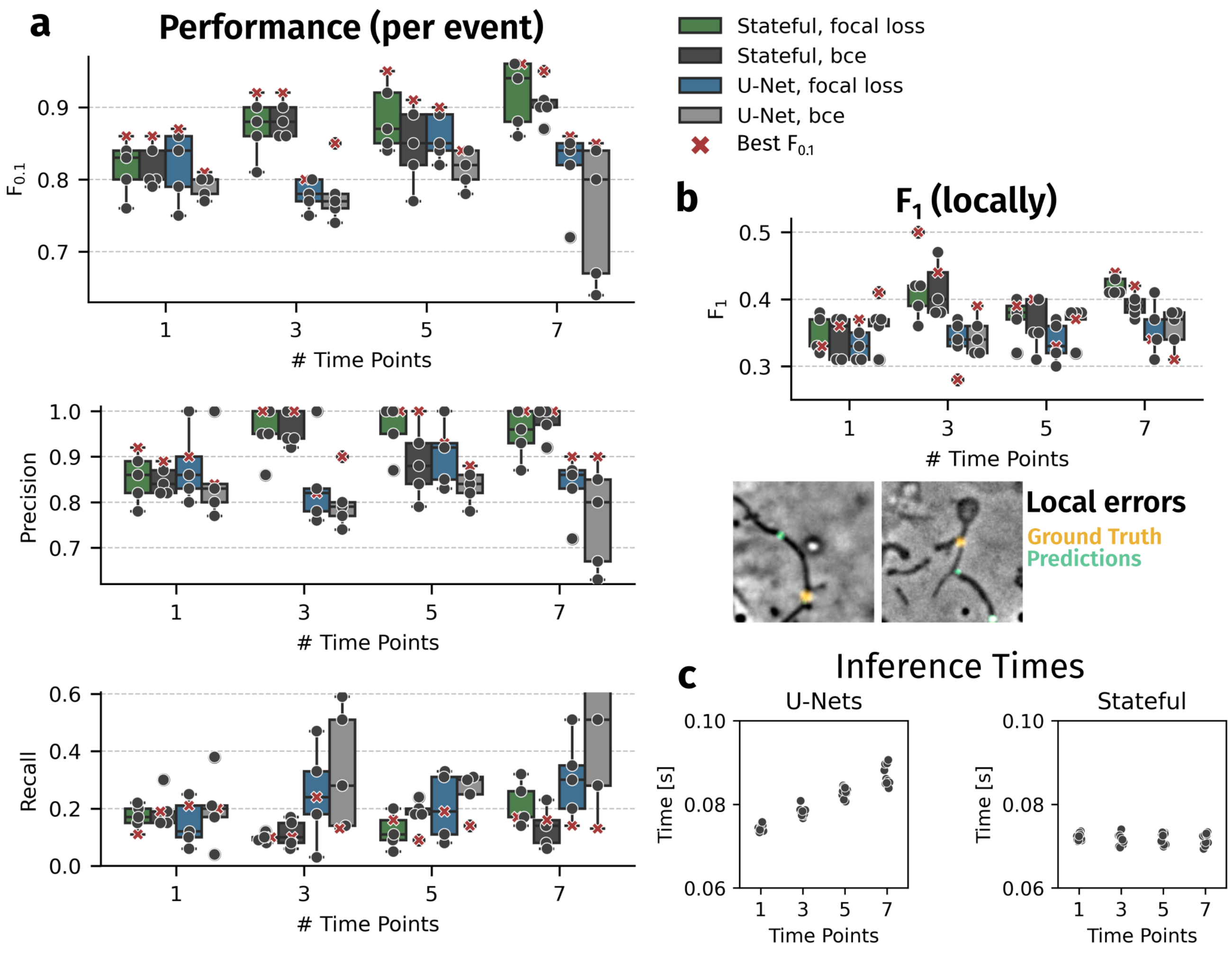


**Fig. S5 | Figure S5 Neural network training for the detection of division pre-states in mitochondria was performed with the different loss functions and architectures presented in this work. a,** Main performance scores calculated for all on an events basis (detection true/false per 5 frames). **b,** Local $F_1$ scores show improvements with additional time points for all architectures and loss functions. Relatively low scores can be explained by detection of constrictions on a mitochondrion that is prone to divide, but in the wrong position. These local false positives are not detrimental to hybrid-EDA, as a trigger on the correct mitochondrion will still acquire the event of interest. **c,** The prediction time of U-Nets increases slightly with more time points due to more data being handled at the same time. As the state in stateful U-Nets can be saved in the layer itself, every individual inference only needs one frame at a time, leading to a constant inference time for on-the-fly processing of a continuous image stream. Each point indicates the performance of a separate model, with an X indicating the best-performing one. Box plots mark the first quartile, median and third quartile with the whiskers spanning the 5th and 95th percentile.

#### 6 - Additional mitochondria division events

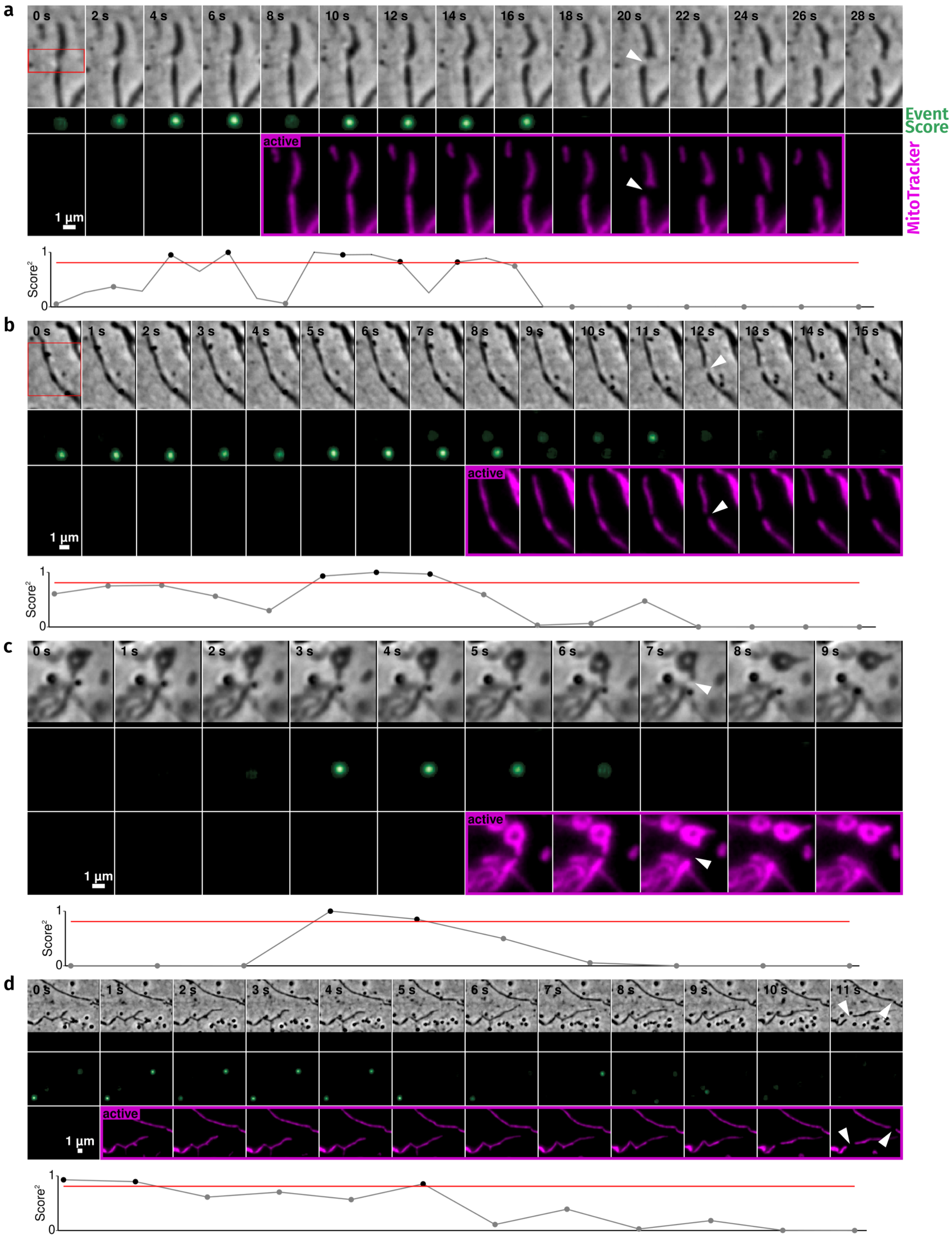


**Fig. S6 | Additional division events captured using hybrid EDA.** A variety of mitochondrial morphologies triggered the adaptive fluorescence acquisitions in different cellular environments. Phase contrast imaging (top), MitoTracker fluorescence (bottom), and score obtained from the model on-the-fly as an image (crop marked as a red square at time 0). The event score is plotted below each series with a red line marking the threshold used.
